# Accumulation of FAM98B and tRNA-Splicing Ligase Complex Proteins in Neuronal Intranuclear Inclusions in Spinocerebellar ataxia type 4

**DOI:** 10.64898/2026.10.26.761951

**Authors:** Mandi Gandelman, Karla P. Figueroa, Anton Classen, Daniel R. Scoles, Stefan M. Pulst

## Abstract

**Background and Objectives:** Spinocerebellar ataxia type 4 (SCA4) is caused by a GGC-repeat expansion in *ZFHX3* encoding an expanded polyglycine tract causing p62- and ubiquitin-positive neuronal intranuclear inclusions (NIIs). In multiple polyglycine diseases NIIs sequester tRNA-splicing ligase complex (tRNA-LC) proteins, causing downstream functional defects linked to neurodegeneration. We assessed the localization of tRNA-LC proteins in SCA4 pontine neurons.

**Methods:** Postmortem brain tissue from a genetically confirmed SCA4 patient was examined across multiple regions using immunofluorescence confocal microscopy for p62, ubiquitin and tRNA-LC proteins FAM98B, RTRAF, and RTCB.

**Results:** In pontine neurons, FAM98B, RTRAF, and RTCB were incorporated into NII, accumulating in a distinct outer layer respective to a ubiquitin inner layer. This redistribution was accompanied by depletion of all three proteins from the soluble nucleoplasm.

**Discussion:** Our findings link SCA4 to a tRNA-LC dysfunction phenotype shared by multiple GGC-repeat disorders, further supporting a unified pathological mechanism in polyglycine diseases.

## Introduction

Spinocerebellar ataxia type 4 (SCA4) is an autosomal dominant, adult-onset cerebellar and sensory ataxia caused by a heterozygous coding GGC-repeat expansion in ZFHX3, also known as ATBF1^1-4^. The expansion encodes an expanded polyglycine tract (polyGly) and is associated with neuronal intranuclear inclusions (NIIs) in basis pontis neurons that are immunoreactive for ZFHX3, p62, and ubiquitin ^1-3^. SCA4 belongs to the expanding group of neurodegenerative GGC/polyGly repeat disorders, but differs from FXTAS and NIID, in which polyGly-containing products arise from repeat-associated translation of noncoding or upstream open reading frames rather than from a coding repeat within a large transcription factor ^5^.

A recently described mechanism in FXTAS and NIID centers on FAM98B, a glycine-rich component of the tRNA-splicing ligase complex (tRNA-LC), which also includes RTRAF/CGI-99, RTCB, DDX1, and Ashwin/ASW ^6^. Through its C-terminal glycine-rich intrinsically disordered region, FAM98B is recruited to polyGly-positive intranuclear inclusions, with associated soluble nucleoplasmic depletion of FAM98B and other tRNA-LC proteins. This depletion disrupts tRNA processing and produces aberrant tRNA-splicing intermediates in patient tissues. In addition, Fam98b depletion in mice caused progressive motor coordination deficits and hindbrain pathology.

We examined histological sections of the pons from a genetically confirmed patient with SCA4 to determine whether FAM98B, RTRAF, and RTCB localize to SCA4 NIIs and whether inclusion-associated redistribution of these proteins is accompanied by soluble nucleoplasmic depletion.

## Methods

### Human subjects

All procedures involving human subjects were approved by the Institutional Review Board (IRB) at the University of Utah. All human subjects gave written consent for inclusion in the study.

### Staining of brain sections

Formalin-fixed, paraffin-embedded pontine sections were deparaffinized in xylene, and rehydrated through graded ethanol to water.

For immunofluorescence, slides underwent heat-mediated antigen retrieval for 10 min in citric acid-based antigen unmasking solution (Vector Laboratories, H-3300), followed by permeabilization in PBS containing 5% normal goat serum and 0.3% Triton X-100 for 30 min. Lipofuscin autofluorescence was quenched with TrueBlack Lipofuscin Autofluorescence Quencher (Biotium) for 60 s, and sections were blocked for 1 h in PBS containing 5% normal goat serum. Primary antibodies were diluted 1:100 in blocking solution and incubated overnight: anti-SQSTM1/p62 (Thermo Fisher Scientific, PA5-20839), anti-ubiquitin (Abcam, ab134953), anti-FAM98B (Proteintech, 22251-1), anti-RTCB (Proteintech, 19809-1), and anti-RTRAF (Proteintech, 19848-1). Slides were incubated with Alexa Fluor Plus 594- and 488-conjugated secondary antibodies (Thermo Fisher Scientific; 1:1,000) for 2 h, counterstained with DAPI, and mounted with ProLong Gold Antifade Mountant (Thermo Fisher Scientific). Confocal images were acquired using a Leica SP8 microscope at the University of Utah Cell Imaging Core.

### Image acquisition and analysis

Images were acquired with a Leica SP8 Confocal Microscope with an oil immersion microscope objective HC PL APO 63x/1,40 OIL CS2. Diffraction-limited 3D z-stack images of pontine neurons were acquired with 0.7AU pinhole size, zoom 4 and 512x512 pixel grid to achieve 90nm pixel size in xy dimension, and with 250nm z-stack spacing. Raw data was further deconvolved with the Leica Lightning software module to achieve best resolution. Images were visualized with LASX (Leica,version 3.5.7). Figure assembly, NII quantification, and fluorescence intensity measurements were performed using ImageJ/Fiji (version 1.54p), including the Plot Profile tool and QuickFigures plugin (version 2023.2). Orthogonal projections and three-dimensional reconstructions were generated using Imaris (Oxford Instruments, version 11.0.1).

For each antibody, images were acquired from at least 10 randomly selected fields containing a minimum of three neurons each. Neurons were defined as cells with nuclear diameters >9 µm. Neuronal intranuclear inclusions >1 µm in diameter were quantified in more than 100 neurons per antibody.

Statistical analyses and data visualization were performed using GraphPad Prism (version 10). Comparisons between groups were made using unpaired Student’s t-tests.

## Results

The tissue used in this study was obtained from a previously reported patient with long-duration SCA4^1^. A summary of the neuropathological findings is presented in Table 1. Pontine sections from this case were examined to determine whether proteins implicated in polyGly-associated tRNA ligase complex (tRNA-LC) pathology localize to neuronal intranuclear inclusions (NIIs), identified by p62 or Ubiquitin immunoreactivity. FAM98B was incorporated into nearly all NIIs examined, with 77 of 80 NIIs immunoreactive for FAM98B (Supplemental Figure 1A). The tRNA-LC proteins RTRAF and RTCB were also detected in NIIs, with 74 of 90 NIIs immunoreactive for RTRAF and 70 of 74 immunoreactive for RTCB (Supplemental Figure 1B-C). Thus, SCA4 NIIs contain FAM98B and multiple tRNA-LC proteins.

**Table 1.**
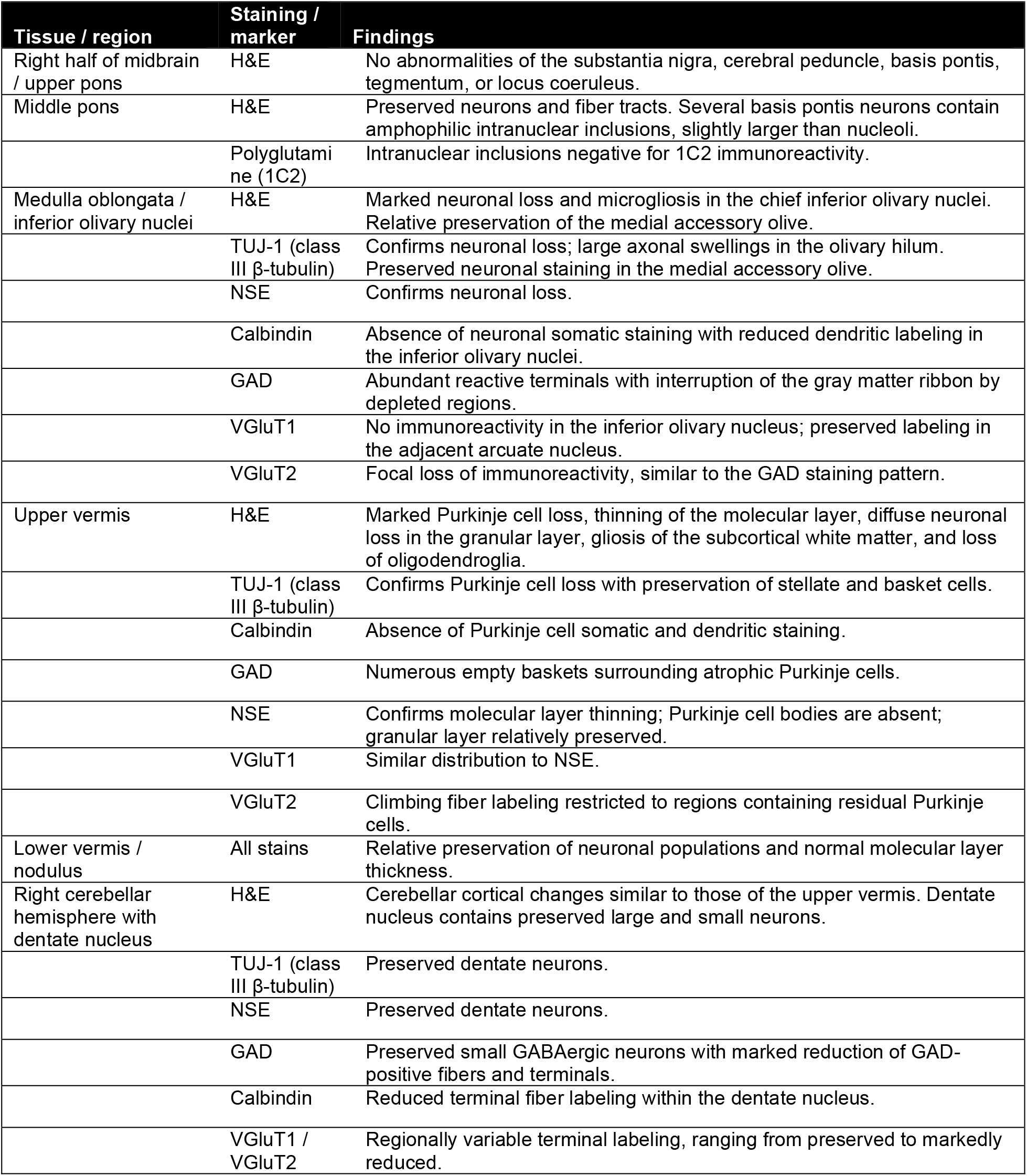
Summary of neuropathological findings in the SCA4 autopsy brain. Neuropathological findings are summarized by brain region and histological or immunohistochemical stain. H&E, hematoxylin and eosin; 1C2, anti-polyglutamine; TUJ-1, class III β-tubulin; NSE, neuron-specific enolase; GAD, glutamate decarboxylase; VGluT1, vesicular glutamate transporter 1; VGluT2, vesicular glutamate transporter 2.

| Tissue / region | Staining / marker | Findings |
| --- | --- | --- |
| Right half of midbrain / upper pons | H&E | No abnormalities of the substantia nigra, cerebral peduncle, basis pontis, tegmentum, or locus coeruleus. |
| Middle pons | H&E | Preserved neurons and fiber tracts. Several basis pontis neurons contain amphophilic intranuclear inclusions, slightly larger than nucleoli. |
|  | Polyglutamine (1C2) | Intranuclear inclusions negative for 1C2 immunoreactivity. |
| Medulla oblongata / inferior olivary nuclei | H&E | Marked neuronal loss and microgliosis in the chief inferior olivary nuclei. Relative preservation of the medial accessory olive. |
| | TUJ-1 (class III β-tubulin) | Confirms neuronal loss; large axonal swellings in the olivary hilum. Preserved neuronal staining in the medial accessory olive. |
|  | NSE | Confirms neuronal loss. |
|  | Calbindin | Absence of neuronal somatic staining with reduced dendritic labeling in the inferior olivary nuclei. |
|  | GAD | Abundant reactive terminals with interruption of the gray matter ribbon by depleted regions. |
|  | VGluT1 | No immunoreactivity in the inferior olivary nucleus; preserved labeling in the adjacent arcuate nucleus. |
|  | VGluT2 | Focal loss of immunoreactivity, similar to the GAD staining pattern. |
| Upper vermis | H&E | Marked Purkinje cell loss, thinning of the molecular layer, diffuse neuronal loss in the granular layer, gliosis of the subcortical white matter, and loss of oligodendroglia. |
| | TUJ-1 (class III β-tubulin) | Confirms Purkinje cell loss with preservation of stellate and basket cells. |
|  | Calbindin | Absence of Purkinje cell somatic and dendritic staining. |
|  | GAD | Numerous empty baskets surrounding atrophic Purkinje cells. |
|  | NSE | Confirms molecular layer thinning; Purkinje cell bodies are absent; granular layer relatively preserved. |
|  | VGluT1 | Similar distribution to NSE. |
|  | VGluT2 | Climbing fiber labeling restricted to regions containing residual Purkinje cells. |
| Lower vermis / nodulus | All stains | Relative preservation of neuronal populations and normal molecular layer thickness. |
| Right cerebellar hemisphere with dentate nucleus | H&E | Cerebellar cortical changes similar to those of the upper vermis. Dentate nucleus contains preserved large and small neurons. |
| | TUJ-1 (class III β-tubulin) | Preserved dentate neurons. |
|  | NSE | Preserved dentate neurons. |
|  | GAD | Preserved small GABAergic neurons with marked reduction of GAD-positive fibers and terminals. |
|  | Calbindin | Reduced terminal fiber labeling within the dentate nucleus. |
|  | VGluT1 / VGluT2 | Regionally variable terminal labeling, ranging from preserved to markedly reduced. |

Imaging of NII in z-stacks and 3D reconstruction showed nucleus contain between 0-4 NIIs in a wide range of size and morphology. While most NII were small to medium spheres (diameter 1-6 µm), some NIIs exhibited complex 3D structures comprised of multiple spherical lobules and occupied over half of the nuclear volume (Figure 1A). Orthogonal projections (Figure 1B) and plot profiles (Figure 1C) reveal a layered organization of tRNA-LC proteins and Ubiquitin. tRNA-LC proteins localized to the outer shell layer of the NII, followed by an area of overlap with Ubiquitin, and an inner shell layer that contained only Ubiquitin immunoreactivity.

**Fig. 1.**
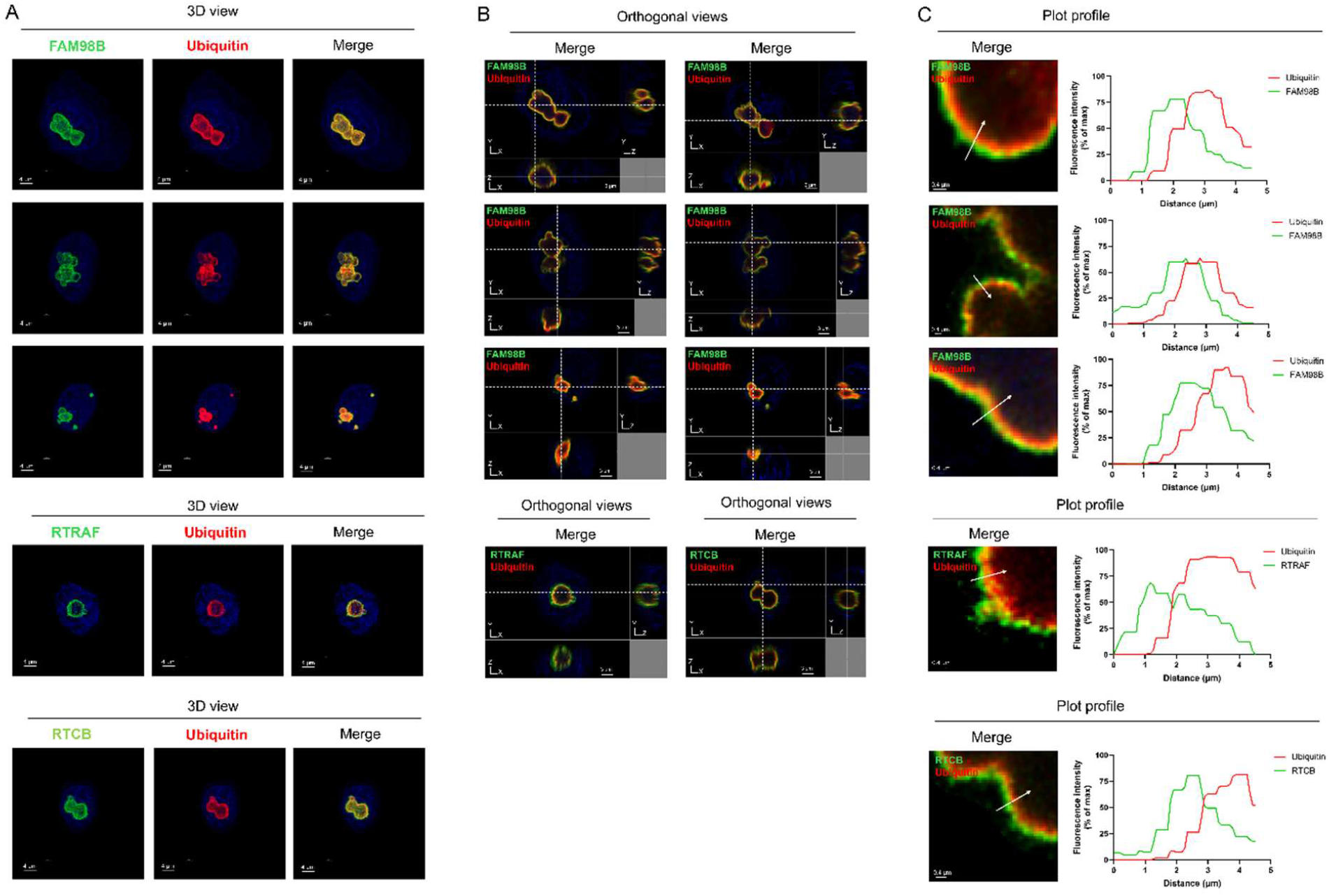
tRNA-LC proteins localize to NII in SCA4 pons. Confocal immunofluorescence microscopy of pontine neurons from an individual with SCA4 stained for FAM98B and ubiquitin (top), RTRAF and ubiquitin (middle), or RTCB and ubiquitin (bottom). (A) Three-dimensional reconstructions of representative NIIs. (B) Orthogonal projections showing the distribution of immunoreactivity in the x–z and y–z planes. (C) Enlarged views of representative NIIs with fluorescence intensity profiles measured along the white arrows, illustrating the relative distribution of tRNA-LC proteins and ubiquitin across the inclusion. Scale bars, as indicated.

Accumulation of tRNA-LC proteins in NIIs has been shown to deplete their soluble nucleoplasmic pool, causing pathological loss-of-function ^6^. Pontine neurons from a control non-SCA4 patient showed uniform nuclear distribution of tRNA-LC proteins (Supplemental Figure 2). In SCA4 pons, neuronal nuclei lacking NII show uniform distribution of tRNA-LC protein immunoreactivity (Figure 2A). In contrast, in neurons containing NII the immunoreactivity was focally enriched in the NII and depleted from the surrounding nucleoplasm (Figure 2A). Quantification of the tRNA-LC protein intensity in the soluble nucleoplasm revealed in this area FAM98B was reduced by 86%, RTRAF by 46% and RTCB by 59% (Figure 2B).

**Fig. 2.**
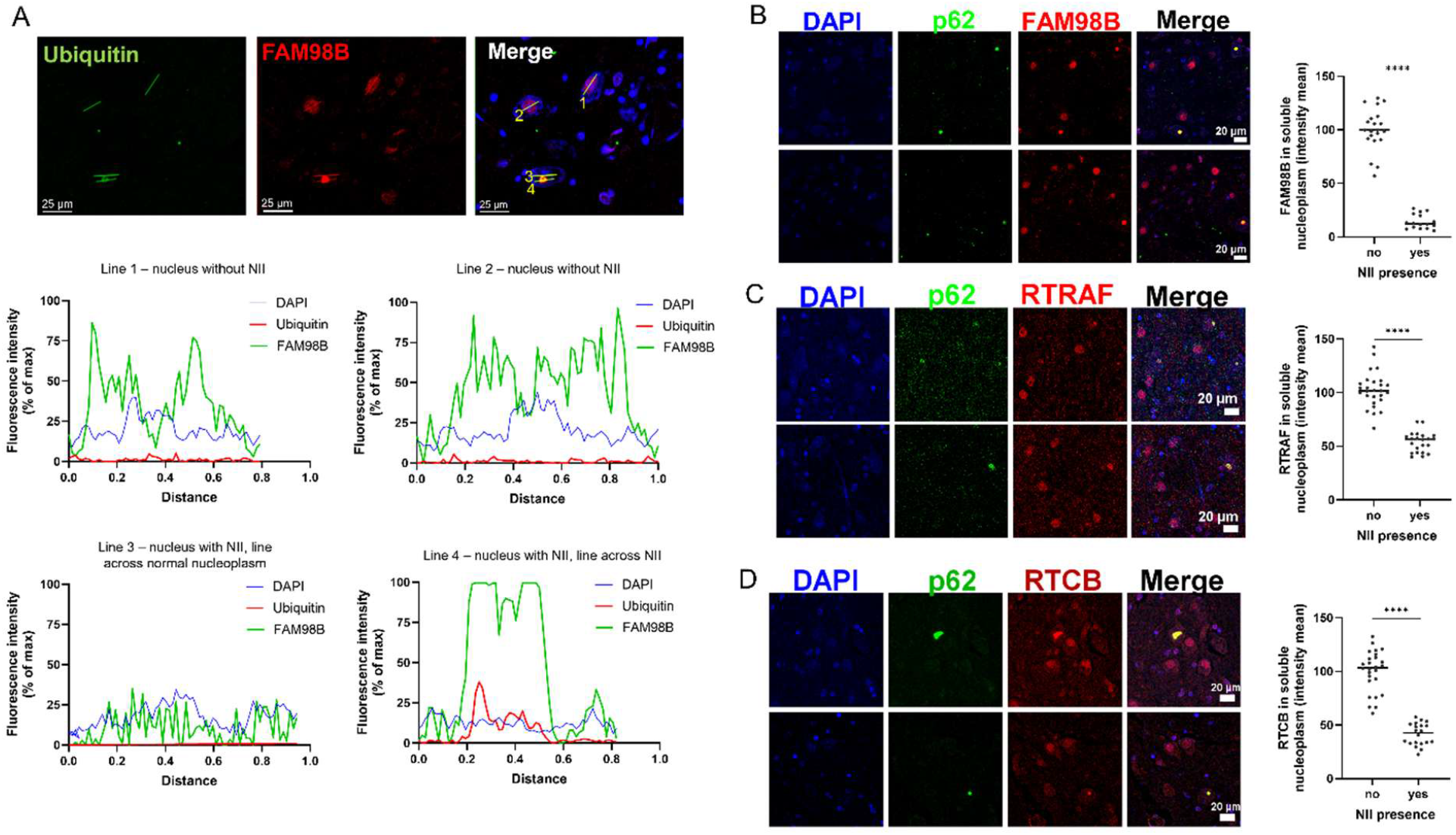
tRNA-LC proteins are enriched within NII and depleted from the surrounding nucleoplasm in SCA4 pons. (A) Representative fluorescence intensity profiles measured along the indicated lines across pontine neuronal nuclei with or without NIIs immunolabeled for FAM98B and ubiquitin. Line profiles illustrate the distribution of FAM98B across the nucleoplasm and within NIIs. (B–D) Confocal immunofluorescence microscopy of pontine neurons immunolabeled for FAM98B and p62 (B), RTRAF and p62 (C), or RTCB and p62 (D), together with quantification of soluble nucleoplasmic fluorescence intensity in neurons with or without NIIs. Fluorescence measurements were obtained from nucleoplasmic regions excluding the inclusion. Each point represents one neuronal nucleus; bars indicate mean ± SD. Statistical comparisons were performed using unpaired *t*-tests. Scale bars, as indicated.

## Discussion

In this study we show that the tRNA ligase complex (tRNA-LC) proteins FAM98B, RTRAF, and RTCB accumulate in NIIs in the SCA4 pons, with a corresponding depletion from the surrounding nucleoplasm. These findings expand the molecular pathology of SCA4 beyond the previously ZFHX3-, p62-, and ubiquitin-positive NII phenotype described in patient tissues and the autophagy defects reported in patient-derived cells^1-3^.

Three-dimensional reconstruction and orthogonal slice views showed that SCA4 NIIs varied considerably in size and morphology, ranging from small spherical structures to large multilobulated aggregates occupying a substantial fraction of the nucleus. Within the NII shell, tRNA-LC proteins were organized as an outer layer, whereas ubiquitin comprised the inner layer, with a distinct zone of overlap between them. The biological significance of NII size, architecture, and layered spatial organization is unknown but may provide insight into the mechanisms of NII formation, their effects on nuclear homeostasis, and their contribution to neuronal dysfunction.

SCA4 is caused by a GGC repeat expansion in *ZFHX3* that encodes a polyGly tract, promoting aggregation of the mutant protein within ubiquitin- and p62-positive NIIs. In polyGly diseases, NIIs recruit endogenous glycine-rich proteins, including FAM98B, whose intrinsically disordered C-terminal region contains the highest glycine content in the human proteome ^6^. In FXTAS and NIID, sequestration of FAM98B is accompanied by recruitment of the remaining tRNA-LC components and depletion of the complex from the soluble nucleoplasm, resulting in impaired tRNA splicing and accumulation of aberrant tRNA-splicing intermediates. Moreover, conditional deletion of Fam98b in adult mice produces motor coordination deficits and selective hindbrain pathology, supporting the functional importance of this pathway^6^.

More broadly, increasing evidence implicates defects in tRNA processing as a mechanism of neurodegeneration. Pathogenic variants in the TSEN tRNA-splicing endonuclease complex, its associated kinase CLP1, and genes involved in tRNA maturation, aminoacylation, or selenocysteine synthesis all cause forms of pontocerebellar hypoplasia or related cerebellar disorders^7-12^. Likewise, disruption of specific unspliced tRNAs in mice results in cerebellar degeneration and ataxia^13,14^.

Our findings extend the tRNA-LC pathology described in FXTAS and NIID to SCA4, suggesting that tRNA-LC dysfunction may represent a convergent feature of polyGly-mediated neurodegeneration. Although the present study is limited to a single autopsy case, the observed redistribution of all three tRNA-LC proteins closely parallels that previously described in FXTAS and NIID, supporting the hypothesis that disruption of tRNA processing is a convergent mechanism underlying polyGly-mediated neurodegeneration.

## Acknowledgments

The authors wish to thank the patient, family, and tissue donors whose participation made this study possible. We acknowledge the Health Sciences Imaging Core at the University of Utah for access to the Leica SP8 confocal microscope and Imaris software. We are grateful to Dr. Arnulf H. Koeppen, Professor of Neurology and Pathology at Albany Medical College and the Albany Stratton VA Medical Center, for providing the neuropathologic evaluation of the SCA4 patient, and to Dr. Frederic C. Clayton for assistance with the autopsy and tissue collection. This work was supported by the National Institutes of Health under award number R33NS124965, R35NS127253, R21NS145030, R01NS137233 and R21NS128630.

This manuscript is the result of funding in whole or in part by the National Institutes of Health (NIH). It is subject to the NIH Public Access Policy. Through acceptance of this federal funding, NIH has been given a right to make this manuscript publicly available in PubMed Central upon the Official Date of Publication, as defined by NIH.” The research was supported by grants and to SMP was additionally supported by the Dale R. Rindlisbacher Endowment Fund and DRS by the John P. Punke Endowment Fund.

---

**Supplementary Fig. 1.**
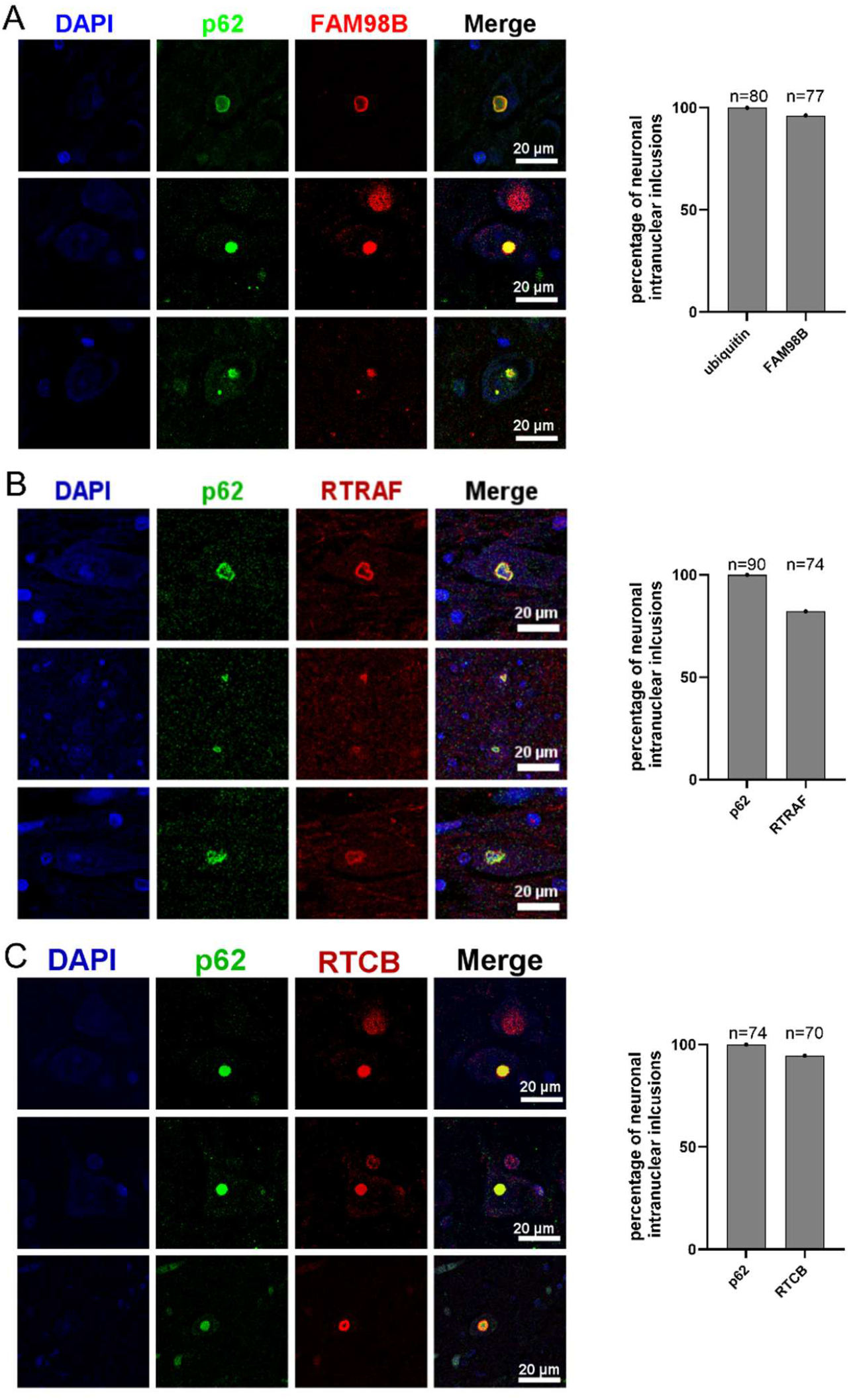
Confocal immunofluorescence analysis of FAM98B, RTRAF, and RTCB localization to neuronal intranuclear inclusions in the SCA4 pons. Representative images of confocal immunofluorescence microscopy and quantification of the percentage of ubiquitin or p62 positive NIIs also immunoreactive for FAM98B (A), RTRAF (B), or RTCB (C) in pontine neurons from an individual with SCA4. Numbers above bars indicate the total number of NIIs in each group. Scale bars, as indicated.

**Supplementary Figure 2.**
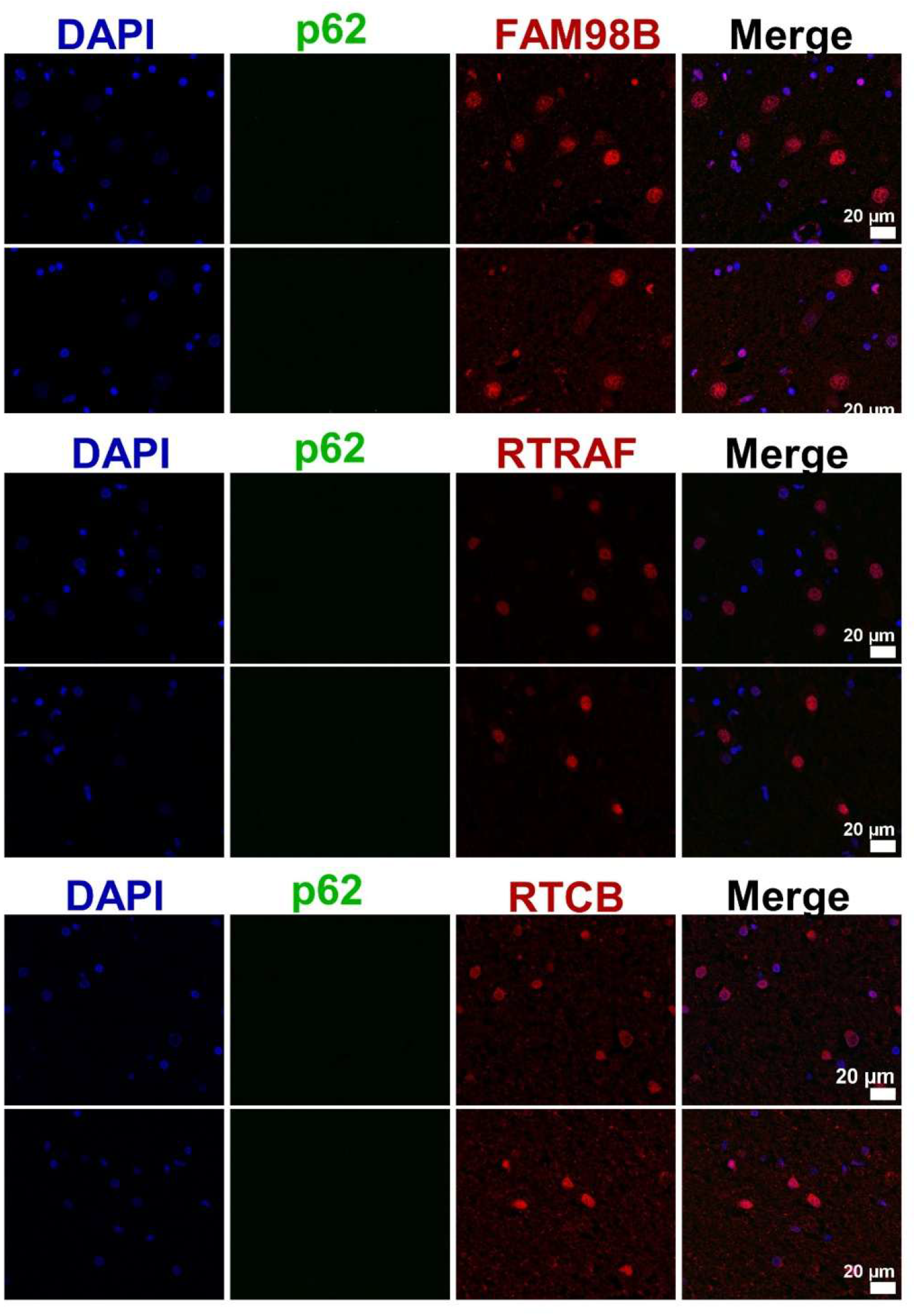
Non-SCA4 control subject basis pontis neurons lack p62-positive NII and show uniform nucleoplasmic distribution of FAM98B and tRNA-splicing ligase complex proteins. Representative immunofluorescence microscopy images of non-SCA4 control basis pontis stained with DAPI, p62, and A, FAM98B; B, RTRAF; or C, RTCB. Scale bars, 20 μm.

## References

1. Figueroa KP, Gross C, Buena-Atienza E, et al. A GGC-repeat expansion in ZFHX3 encoding polyglycine causes spinocerebellar ataxia type 4 and impairs autophagy. Nat Genet. Jun 2024;56(6):1080–1089. doi:10.1038/s41588-024-01719-5

2. Wallenius J, Kafantari E, Jhaveri E, et al. Exonic trinucleotide repeat expansions in ZFHX3 cause spinocerebellar ataxia type 4: A poly-glycine disease. Am J Hum Genet. Jan 4 2024;111(1):82–95. doi:10.1016/j.ajhg.2023.11.008

3. Paucar M, Nilsson D, Engvall M, et al. Spinocerebellar ataxia type 4 is caused by a GGC expansion in the ZFHX3 gene and is associated with prominent dysautonomia and motor neuron signs. J Intern Med. Sep 2024;296(3):234–248. doi:10.1111/joim.13815

4. Chen Z, Gustavsson EK, Macpherson H, et al. Adaptive Long-Read Sequencing Reveals GGC Repeat Expansion in ZFHX3 Associated with Spinocerebellar Ataxia Type 4. Mov Disord. Mar 2024;39(3):486–497. doi:10.1002/mds.29704

5. Liufu T, Zheng Y, Yu J, et al. The polyG diseases: a new disease entity. Acta Neuropathol Commun. May 31 2022;10(1):79. doi:10.1186/s40478-022-01383-y

6. Yang J, Xu Y, Ziehr DR, et al. Polyglycine-mediated aggregation of FAM98B disrupts tRNA processing in GGC repeat disorders. Science. Jul 17 2025;389(6757):eado2403. doi:10.1126/science.ado2403

7. Budde BS, Namavar Y, Barth PG, et al. tRNA splicing endonuclease mutations cause pontocerebellar hypoplasia. Nat Genet. Sep 2008;40(9):1113–8. doi:10.1038/ng.204

8. Edvardson S, Shaag A, Kolesnikova O, et al. Deleterious mutation in the mitochondrial arginyl-transfer RNA synthetase gene is associated with pontocerebellar hypoplasia. Am J Hum Genet. Oct 2007;81(4):857–62. doi:10.1086/521227

9. Schaffer AE, Eggens VR, Caglayan AO, et al. CLP1 founder mutation links tRNA splicing and maturation to cerebellar development and neurodegeneration. Cell. Apr 24 2014;157(3):651–63. doi:10.1016/j.cell.2014.03.049

10. Karaca E, Weitzer S, Pehlivan D, et al. Human CLP1 mutations alter tRNA biogenesis, affecting both peripheral and central nervous system function. Cell. Apr 24 2014;157(3):636–50. doi:10.1016/j.cell.2014.02.058

11. Agamy O, Ben Zeev B, Lev D, et al. Mutations disrupting selenocysteine formation cause progressive cerebello-cerebral atrophy. Am J Hum Genet. Oct 8 2010;87(4):538–44. doi:10.1016/j.ajhg.2010.09.007

12. Breuss MW, Sultan T, James KN, et al. Autosomal-Recessive Mutations in the tRNA Splicing Endonuclease Subunit TSEN15 Cause Pontocerebellar Hypoplasia and Progressive Microcephaly. Am J Hum Genet. Sep 1 2016;99(3):785. doi:10.1016/j.ajhg.2016.08.009

13. Ishimura R, Nagy G, Dotu I, et al. RNA function. Ribosome stalling induced by mutation of a CNS-specific tRNA causes neurodegeneration. Science. Jul 25 2014;345(6195):455–9. doi:10.1126/science.1249749

14. Wirth EK, Bharathi BS, Hatfield D, Conrad M, Brielmeier M, Schweizer U. Cerebellar hypoplasia in mice lacking selenoprotein biosynthesis in neurons. Biol Trace Elem Res. May 2014;158(2):203–10. doi:10.1007/s12011-014-9920-z

